# Still moving: The double-drift illusion survives smooth pursuit

**DOI:** 10.1101/624296

**Authors:** Patrick Cavanagh, Peter U. Tse

**Affiliations:** Department of Psychology, Glendon College, CVR York University, Toronto, ON M4N3M6, Canada; Department of Psychological and Brain Sciences, Dartmouth College, Hanover, NJ 03755, USA

**Keywords:** Motion, motion-induced position shift, smooth pursuit

## Abstract

If a gabor pattern drifts in one direction while its internal texture drifts in the orthogonal direction, observers see a remarkable shift in its perceived direction when it is viewed in the periphery. The reported direction of the double-drift stimulus (also known as the infinite regress and curveball illusions) is some combination of the actual external motion of the gabor envelope and the internal motion of its texture (Tse & Hsieh, 2006). Here we find that if the observers track a fixation point that moves in tandem with the gabor, the illusion is undiminished. The pursuit of the moving fixation spot keeps the gabor roughly fixed at one location on the retina, cancelling its external motion, leaving only the internal motion. The gabor is seen to move in the world at roughly its actual speed as the motion of the eye is discounted at some point to recover velocities in world coordinates (e.g. Wallach, 1959). Our finding indicates that the combination of internal and external motion that produces the double drift illusion must happen after the eye movement signals have been factored into stimulus motions. We also test the double drift effect at various path lengths, durations, and speeds, with both mid-grey and black backgrounds, all with a static fixation. These results confirm that a simple vector combination of the two speeds alone accounts for virtually all the direction shifts on the grey background. On the black background, the illusion is eliminated. These results place constraints on where perceived spatial coordinates arise in the visual processing hierarchy to locations at or beyond where compensation for pursuit eye movements arise, specifically V3A, V6, MSTd, and VIP (e.g., Nau et al, 2018).

## Introduction

Most cells in the visual system are characterized by receptive fields, organized in retinotopic maps. The location specificity of a receptive field provides a straightforward positional coding of any stimulus that activates the cell. It might seem that there is little further to be discovered concerning the computation of position, but that would be wrong. Dramatic errors in perceived location are found when the target (Nijhawan, 1994; Eagleman & Sejnowski, 2007; Cavanagh & Anstis, 2013) or the eyes (review Ross et al, 2001) are in motion. These results indicate that perceived location must instead be computed by a sophisticated, predictive system that constructs target locations based on, most importantly, their motion. We show here that this system lies at or beyond the level at which eye movements are discounted to recover motions in the world.

One particular stimulus, the double-drift stimulus (Tse & Hsieh, 2006; Shapiro et al, 2010; Gurnsey and Biard, 2012; Kwon et al, 2015; Lisi & Cavanagh, 2015) leads to extreme misjudgments of an object’s location (Fig. 1, left). In this stimulus, a gabor patch moves over an equiluminant background in one direction while its internal texture moves orthogonally to the gabor’s path (Fig. 1). The perceived direction of this double-drift stimulus (Lisi & Cavanagh, 2015) may differ by more than 50° in direction from its physical direction. Moreover, the effect is also seen for the judged position of the target (Lisi & Cavanagh 2015) which may deviate by as much as several degrees of visual angle from its true location.

**Figure 1.**
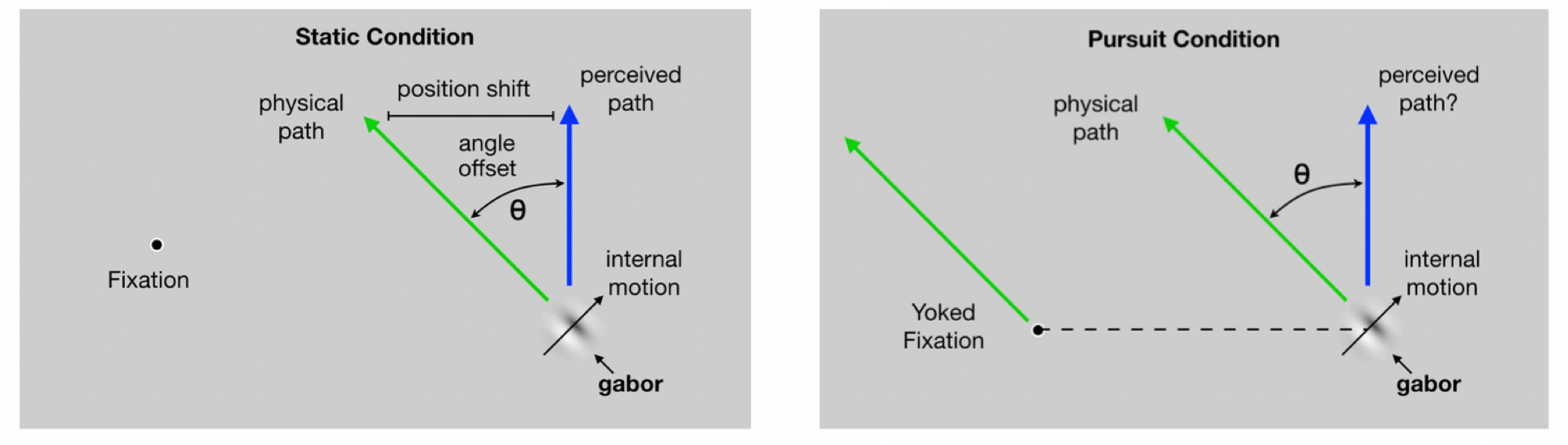
**Left.** A large shift in perceived direction and perceived position is produced when the internal texture of a moving gabor drifts orthogonally to the gabor’s path. **Right.** If the fixation point moves in tandem with the gabor, then the position of the gabor on the retina is roughly stable. Here we test if the illusion survives the rough stabilization of retinal motion that arises during this pursuit condition. See Movie 1 and Movie 2 (at the moment, this only down loads the two movies, I will set up a web page to display them when you click).

Surprisingly, this robust perceptual illusion does not affect saccadic eye movements to the gabor which are executed to their physical not their perceived location (Lisi & Cavanagh, 2015). This suggests that saccade control areas, such as the superior colliculus or frontal eye fields, program a trajectory to where the gabor is, not to where it appears to be, as long as the gabor is present while the saccade is being programmed. If the saccade is programmed after the gabor has been removed from the display (Massendari et al, 2018) the illusion does affect saccade landings. Presumably, when the gabor is not present in the stimulus, the saccade computation must rely on the memory of its location, implying that the memory trace used by the saccade system is in perceptual coordinates. Similarly, pointing responses (Lisi & Cavanagh, 2017) to the double drift stimulus show the illusion, indicating that pointing is also governed by the perceived location.

Here we examine whether the illusion persists when the participants smoothly pursue a fixation point that is yoked to the motion of the gabor (Fig. 1, right). With roughly accurate pursuit, the gabor will have little motion on the retina even though its motion in the world is clearly seen. Clearly, the visual system can recover motion in the world during pursuit by compensating for the motion of the eye either based on retinal or extra-retinal inputs (Wallach, 1959; Rieger & Lawton, 1985; Brenner & van den Berg, 1996; Freeman, 2001; Souman, Hooge, & Wertheim, 2006; Freeman, Champion, Sumnall, & Snowden, 2009). If the illusion fails during smooth pursuit (as it did for saccades, Lisi & Cavanagh, 2015), then the motion computation that produces the illusory direction must occur prior to the correction for eye movement. If the illusion is maintained during pursuit, the vector combination must be at or after the point at which motions in the world have been recovered. We also measure the double-drift illusion at various speeds of the gabor and on a black background to examine how the internal and external motion vectors are combined to produce the perceived direction.

## Experiment 1. Eyes fixed vs smooth pursuit

This experiment measured the perceived shift in the orientation of the gabor’s path when the fixation was either static or moving. The gabor moved continuously back and forth along an oblique path with its internal motion reversing direction at each endpoint so the apparent orientation of the path appeared stable (Lisi & Cavanagh, 2015). Participants adjusted the orientation of the gabor’s path until it appeared vertical. When the fixation point was moving, it followed the path of the gabor, offset horizontally to the left by 18.75 degrees of visual angle (dva) so that the gabor would fall more or less at 18.75 dva to the right of the fovea during the smooth pursuit.

## Method

### Participants

Six healthy adults took part in the experiment (2 males, 4 females, mean age = 36.2 years, SD = 22.1, with a range of 16 to 66), including the two authors and four participants naïve to the purpose of the experiment. All participants in this and the following experiment reported normal or corrected-to-normal vision. All participants gave informed consent in writing prior to participation and the protocols for the study were approved by the Dartmouth College Review Board.

### Stimuli and apparatus

The experiment took place in a darkened room. Stimuli were presented on a gamma-corrected CRT monitor (75 Hz, 1152 × 870 resolution covering 36 × 27 cm). Participants were seated 57 cm from the monitor with their heads resting on a chin- and headrest.

The stimulus was a 100% contrast gabor with a sigma of 0.625 degrees of visual angle (dva) and spatial frequency of 1.56 cpd moving at 5 Hz orthogonally to the gabor’s path. This resulted in an internal speed of the gabor’s bars of 7.8 dva/s. In both static fixation and pursuit trials, the gabor moved back and forth over a path 4.69 dva long, reversing direction every 1.0 s for an external speed of 4.69 dva/s. The midpoint of the gabor’s path was 9.375 dva to the right of the center of the screen. The gabor was presented against a mid-gray background (10.2 cd/m^2^). A black (2.1 cd/m^2^) fixation dot remained on the screen throughout the experiment. On the static fixation trials, the fixation point was centered vertically on the screen and 9.375 dva to the left of the screen’s midpoint, so a total of 18.75 dva from the gabor path’s midpoint. On the moving fixation trials, the fixation followed the path of the gabor, offset horizontally 18.75 dva to its left.

### Procedure and design

Participants completed 24 trials, 12 with static fixation and 12 with pursuit. During each trial, the gabor appeared initially at the midpoint of its path. It then moved up to the left, toward its upper endpoint where it reversed both its external and internal motion directions and returned through the midpoint to the bottom extent of its path where it again reversed internal and external directions. These back-and-forth transits repeated throughout the trial. Participants were instructed to keep their gaze fixed on the black dot whether it was stationary or moving and to adjust the orientation of the gabor’s path, using a track pad, until it appeared vertical. When the participant was satisfied with the setting, he or she pressed the space bar to end the trial. The starting orientation of the gabor was set randomly on each trial. The static and pursuit trials were randomly intermixed.

## Results

The magnitude of the illusion was reported as the angular offset from vertical required to make the path appear vertical. Figure 2 shows the results where the average value of the illusion across the 6 participants was 53.45° ± 2.46° for the static fixation condition and 55.54° ± 3.54° for the smooth pursuit condition. A t-test showed no significant difference between the two conditions.

**Figure 2.**
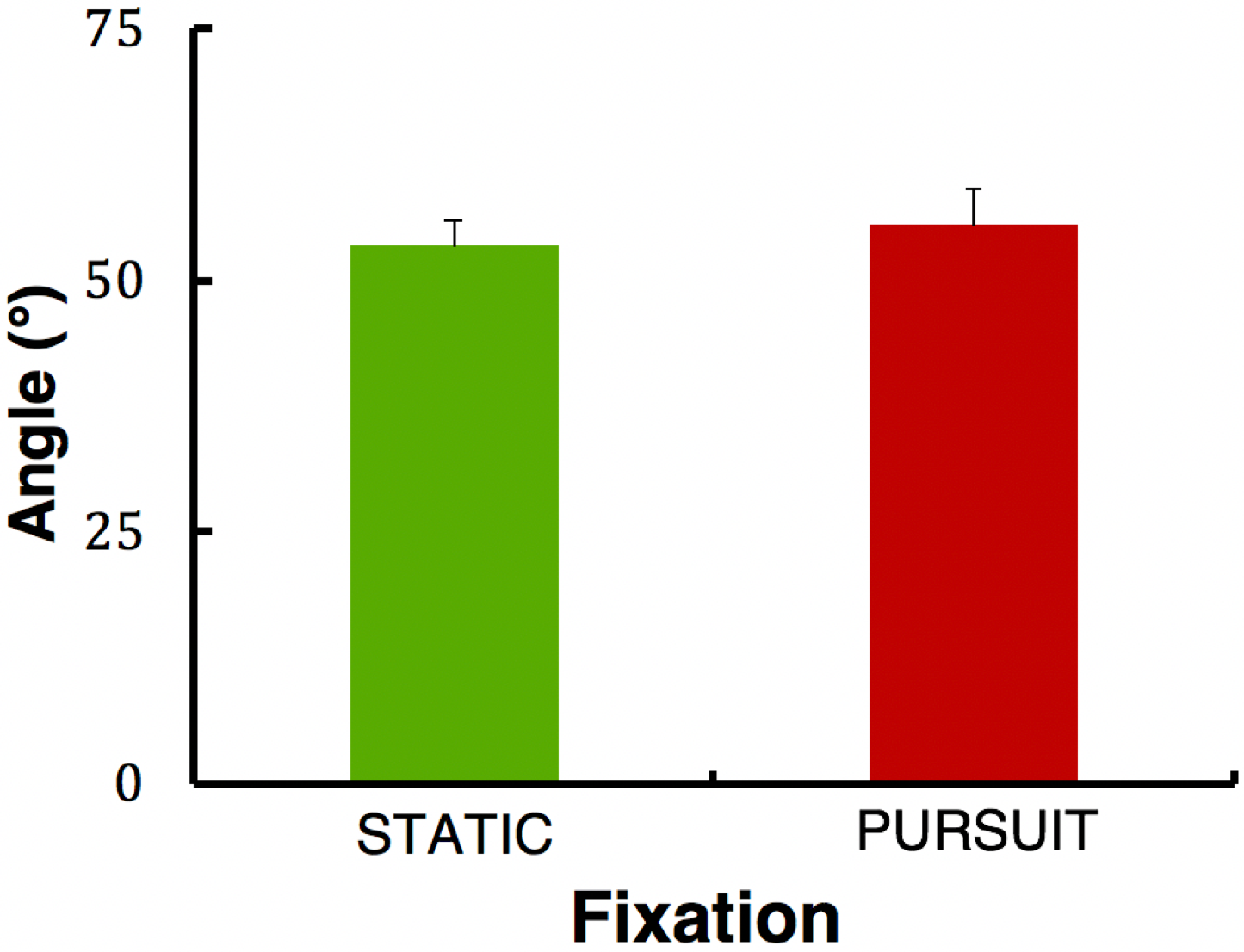
The angular offset from vertical that made a double-drift trajectory appear vertical in the static fixation (in green) and smooth pursuit (in red) conditions. The small bars indicate +1.0 SE.

Figure 2 shows that the double drift illusion clearly survives concurrent smooth pursuit that roughly stabilizes the gabor on the retina. This suggests that the point at which the two motion vectors are combined must be at or following the point where the pursuit motion is combined with retinal motion to recover motions in the world. In the following experiment, we examine how the two vectors, internal and external, combine.

## Experiment 2. Eyes fixed, changing speed and background

This experiment evaluated the effect of the speed of the gabor along its path on the orientation shift. This was tested with the gabor on a mid-grey background with the same mean luminance as the gabor and on a black background (the internal speed of the gabor was constant throughout). If the perceived direction is some fixed combination the internal and external speeds, the perceived direction should follow a simple function of the two speeds (Fig. 3). We varied the speed in two ways: changing the rate at which the motion reversed (every 200, 400, or 800 ms) and changing the path length (3.125 dva and 6.25 dva). We then ran separate sessions with the same speed variations for a gabor on a black background. As before, participants adjusted the orientation of the gabor’s path until it appeared vertical.

**Figure 3.**
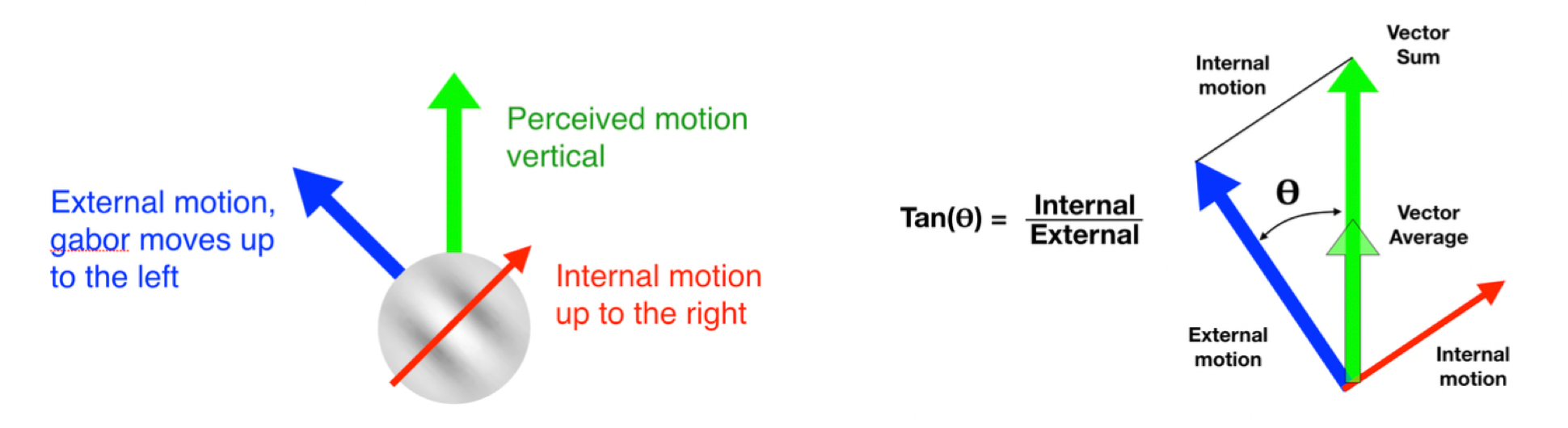
Here we assume that the perceived motion direction is a weighted combination of the internal and external motion vectors. In the experiment, the internal and external vectors are orthogonal so the physical path that appears vertical is given by tan-^1^(internal speed/external speed) when the internal and external speeds are weighted equally, as they would be for a vector sum or a vector average (see Equation 2 in the text).

We can derive the path orientation, **θ**, that will appear vertical assuming that the orthogonal internal and external motion vectors, ***v***_***i***_ and ***v***_***e***_, are combined with weights of ***w***_***i***_ and ***w***_***e***_ (see Fig.3).

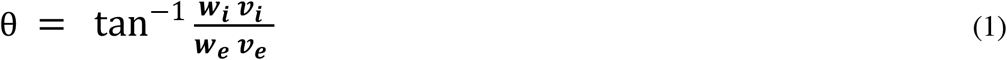

In Figure 3, the weights for the internal and external motions are equal, as they are for a vector sum (***w***_***i***_ = ***w***_***e***_ = 1) or a vector average (***w***_***i***_ = ***w***_***e***_ = 0.5). In our experiments, we evaluate only the perceived angle so we cannot individually estimate the separate weights but only their relative value, ***k*** = ***w***_***i***_/***w***_***e***_. That is what we will recover from our measurements of the angle, **θ**, that appears vertical:

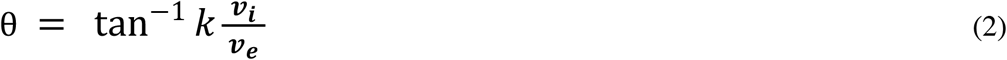

If we also measured the perceived speed, we could recover the individual weights where the speed, the amplitude of the perceptual vector,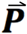, is given by

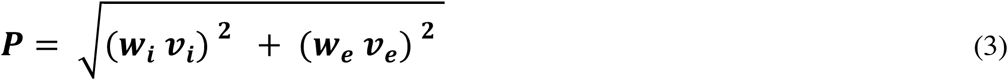

## Method

### Participants

The participants of Experiment 1 all took part in this experiment.

### Stimuli and apparatus

The stimulus and apparatus were the same as in the previous experiment with the following differences. The gabor moved back and forth over one of two path lengths (3.125 dva and 6.25 dva), and reversed direction with one of three durations, 200 ms, 400 ms, or 800 ms. In the main condition, the same gabor from Experiment 1 was again presented against a mid-gray background (10.2 cd/m^2^) that matched its mean luminance. In the black condition, the background was black (2.1 cd/m^2^) and the gabor was 100% contrast with a spatial frequency of 1.56 cpd moving at 5 Hz but it was now in a hard aperture of 1.875 dva diameter with a sigma of 1.875 dva, so it appeared as almost a uniform grating within a circular aperture. The black fixation dot enclosed in a white outline circle remained on the screen throughout the experiment and did not move.

### Procedure and design

Participants performed 24 method of adjustment trials in the main experiment and 24 in the black background experiment, 4 trials in each of the 6 conditions (2 path lengths and 3 reversal rates). The six speed conditions were randomly intermixed. The main and black background experiments were run separately. The procedure was otherwise similar to that of Experiment 1.

## Results

As in Experiment 1, the magnitude of the illusion was reported as the angular offset from vertical required to make the path appear vertical. In Figure 4, the results for the grey background tests are shown in red and blue and reveal that the illusion strength decreased smoothly with the external speed of the gabor (the internal speed was always 5 Hz). Note that the combinations of short and long paths and 3 reversal rates resulted in two pairs of conditions having the same external speeds but different path lengths and durations. Here the red and blue data overlap very closely suggesting that the controlling factor is the external speed and not the path length or reversal rate. We also plot the curve of the direction derived from the internal and external motions (Equation 2) as two dashed lines on the plot, one for a relative weight, ***k***, of 1.0, the weight for a standard vector sum, and one for the relative weight of 0.81 which gives the best fit to the data. The data fall very close to this second line (r^2^ = 0.997) where the internal motion is weighted about 80% of the external motion.

**Figure 4.**
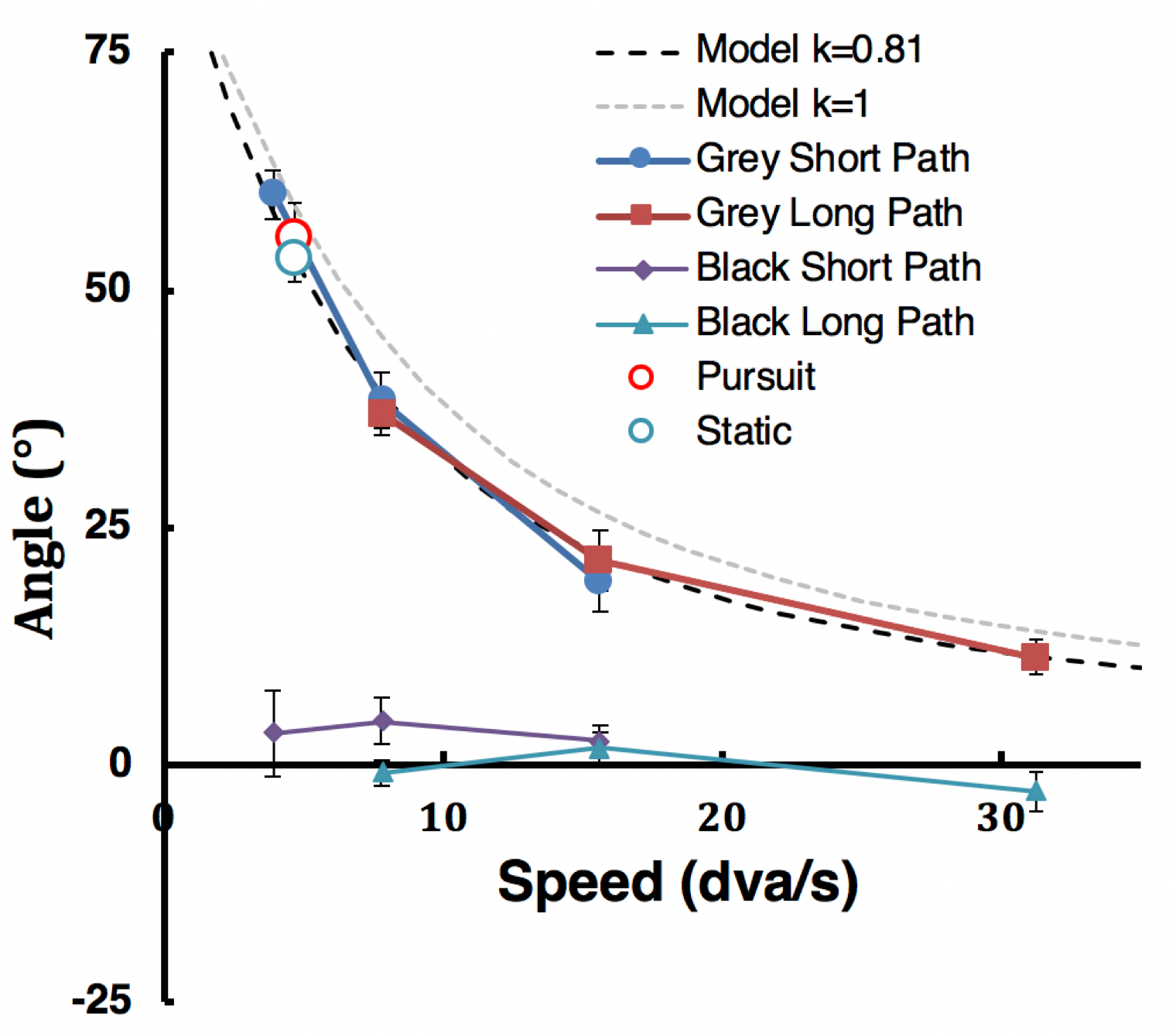
Illusion strength as the angular offset from vertical required to make the path appear vertical. The angle is plotted as a function of the external speed of the gabor and that is given by the path length / reversal time. The model of direction from Equation 3 is given by the two dashed lines, the light one for ***k***=1, equivalent to a vector sum or vector average, the darker one for ***k***=0.81, the best fitting value, indicating that the external motion is given slightly more weight than the internal motion. The vertical bars show ±1 SE.

The results for the black background (green and purple plots on Figure 4) are clear as well – the illusion is significantly weakened or eliminated when the background luminance is no longer matched with the mean luminance of the gabor.

Finally, we replot the two data points from Experiment 1 in Figure 4 (the red and blue outline circles) and they fall on the same line as those with the grey background in this second experiment. Again, this confirms that here with the internal speed fixed, it is the external speed alone that controls the illusion strength and not the factors of path length or reversal rate. The first experiment had a path length (4.69 dva) and reversal time (1000 ms) both differing from the corresponding values in this second experiment. Nevertheless, the resulting external speed places the data on the same function as the new data with static fixation. Thus, the speed that governs the illusion strength must be that of the external motion, specifically in world coordinates, as that is the only frame of reference in which the external speed that is matched for both the static fixation and pursuit conditions. The retinal speed of the external motion is dramatically different in the two conditions being more or less 0 in the pursuit case.

## General discussion

Because the fovea has the highest visual acuity, two types of eye movements have evolved to keep targets of interest at or near the fovea. Saccades bring a peripheral target into the fovea ballistically, while smooth pursuit eye movements keep a target that is moving in the world centered in the fovea (e.g., Bridgeman, Deubel, & Haarmeier, 1999). Lisi and Cavanagh (2015) showed that saccades are made to the stimulus location of the doubly drifting gabor, even when its perceived position may be many degrees of visual angle away from the stimulus location. This suggests that saccades are computed over object positions in retinotopic coordinates, not perceptual coordinates. Here we examined whether the double drift illusion is affected by another class of eye movements: smooth pursuit. We instructed participants to keep fixation on a spot that moved in tandem with the gabor, keeping it roughly fixed on their retina. They then adjusted the perceived path to be vertical. Their settings show that a similar offset from vertical was required to make the path appear vertical whether fixation was static or yoked to the moving gabor. The results show that the double-drift illusion is maintained during smooth pursuit of a moving fixation point that roughly stabilizes the gabor on the retina.

During smooth pursuit, objects that are stationary in the world are displaced across the retina, but this spurious retinal motion is mostly discounted to recover their true motion in world coordinates (Wallach, 1959; Rieger & Lawton, 1985; Brenner & van den Berg, 1996; Freeman, 2001; Souman, et al., 2006; Freeman, et al., 2009). In the case of the double-drift illusion with pursuit of the yoked fixation in Experiment 1, the gabor will be roughly stable on the retina – its average motion will be near zero. Discounting of the pursuit motion will recover, at some level, a perception of the gabor’s true displacement on the display screen. As we discuss in more detail below, the double-drift illusion arises from an inappropriate combination of the gabor’s two motion vectors, the internal and external motion. Experiment 1 suggests that this vector combination occurs along with or after the discounting of the consequences of the smooth pursuit eye movement. If the vector combination occurred before compensating for the pursuit, the gabor envelope would be roughly static on the retina, and we would expect only the much smaller position offset typically seen for the static illusion (De Valois & De Valois, 1991; Ramachandran & Anstis, 1990). Previous research has shown that a static gabor with internal drift is perceived to shift only a small amount, less than the width of the gabor’s aperture (Chung, Patel, Bedell, & Yilmaz, 2007) and this position shift accumulates only over the first 100 ms of the presentation (Chung, et al., 2007). In contrast, the position shift for a moving gabor (the double-drift stimulus) is large, several times the width of its aperture (Lisi & Cavanagh, 2015), and accumulates over the entire duration of the presentation, here up to 1 second. If the vector combination occurs with or after the compensation for the smooth pursuit motion, then the gabor recovers its external motion and the illusion should be similar in magnitude to the double-drift with static fixation. This is what we found.

### Anatomical locus of vector combination

Our results in Experiment 1 showed that smooth pursuit has little effect on the perceived path of the double-drift stimulus. This suggests that the illusion arises at or beyond the neural populations that compensate for self-generated motions, taking retinal slip, image motion, eye and head motion into account to recover object motion in the world. There are several areas with such cells, specifically MST, V3A, V6, and posterior parietal (VIP). Galletti and colleagues originally reported these “real motion” cells in areas V6 and V3A in monkey as well as in early visual cortex (see review Galletti & Fattori, 2003). However, Erickson and Thier (1991) and Thier and Erickson, (1992) reported that the direction-selective cells they found in VI, V2, and V3 showed no compensation for eye movements. They also found that most cells in V4 and MT showed no compensation either. In contrast, cells in dorsal MST did responded preferentially to externally induced motion, suppressing responses to retinal motion caused by eye movements. This finding of cells in MST that responded to motion in the world independently of motion on the retina was supported by several subsequent studies (Dukelow et al., 2001; Ilg & Thier, 2003; Inaba et al 2007). Cells that compensated for eye movements were also reported in parietal areas (Duhamel et al, 1997; Ilg, Schumann, & Thier, 2004).

Results from human fMRI support the results from monkey physiology showing responses to real word motion in V6, V3A and VIP (Schindler & Bartels, 2018; Fischer et al, 2012; Nau et al, 2018). Nau et al. (2018) also report real world motion responses in early areas V1 and V2 but suggested that this was due to feedback from higher areas. Such feedback may also have accounted for Galletti et al.’s (1984) initial reports of “real motion” cells in early visual cortex.

These results indicate that the double-drift illusion must emerge at a later stage than early cortices V1, V2, V3, or MT for the illusion to persist undiminished during pursuit. Our results do not help us differentiate among the several candidate areas that do compensate for eye movements, principally, V3A, V6, MST, and VIP and of course any higher-level areas that share this coding of real world motion. Interestingly, individual cells in these areas can show responses to motion in world coordinates, but across each area, the receptive fields of cells themselves are organized in retinotopic maps for position no matter what the eyes are doing. The motion responses compensate for the retinal consequences of eye movements but the position representation of the cells is not converted into world or spatiotopic coordinates.

Finally, although the physiological and imaging results indicate that cells in these areas do compensate for the motion caused by pursuit eye movements, these findings do not reveal how that compensation comes about. Some authors have suggested that the common motion vector across the retina gives an estimate of the motion created by eye movements (Wallach, 1959; Brenner & van den Berg, 1996; Rieger & Lawton, 1985). Others propose that extra-retinal signals, efference copy, from the motor system would provide the direction and speed of the motion that needs to be discounted (Freeman, 2001; Souman, Hooge, & Wertheim, 2006). Neither the studies reviewed above nor our findings with pursuit here differentiate between these alternatives.

### Incremental accumulation model

The double-drift effect has two components, first, an angle deviation where the gabor’s direction is influenced by its internal motion and second, a continuous accumulation of position offsets where the next perceived position is offset from the current one in the illusory direction. Our experiment has addressed only the angle deviation and we will discuss it first before passing on to the question of the position shifts that accompany the effects on angle.

The angle settings here are consistent with the replacement of the external motion of the gabor with a combination of its internal and external motions as proposed by Tse & Hsieh (2006). The physical angle that makes a double-drift gabor appear move vertically is tan^−1^(***k*** • ***v***_***i***_/***v***_***e***_) where ***k*** is the relative weight of the internal, ***v***_***i***_, and external, ***v***_***e***_, motions in the vector combination (see Equation 2 above and the schematic in Figure 2). This function is plotted twice in Figure 3 along with the data of Experiments 1 and 2, once for equal weights of the internal and external motions (***k*** = 1.0) as would be the case for a standard vector sum or average, and once for the best fitting value of ***k***, 0.81. Tse and Hsieh (2006, Figure 5) found that ***k*** increased non-linearly with increasing ***v***_***i***_, reaching ~0.75 at the highest ***v***_***i***_ tested, 6 dva/s, close to our ***k*** of 0.81 here for a ***v***_***i***_ of 7.69 dva/s. In contrast, Tse and Hsieh found that changes in the external motion, ***v***_***e***_, had no effect on ***k***, and that there was no interaction between ***v***_***i***_ and ***v***_***e***_ on the value of ***k***.

**Figure 5.**
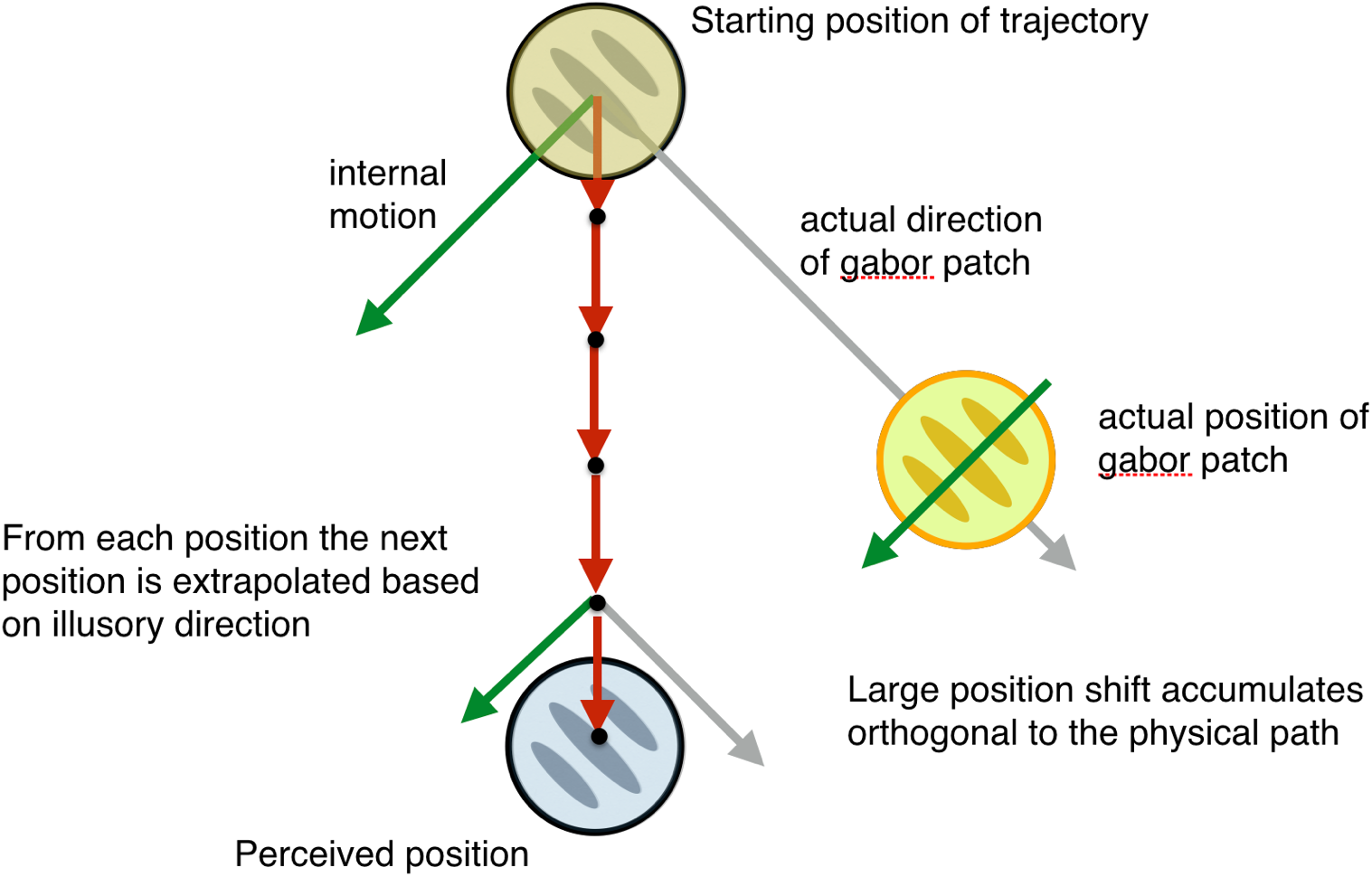
Continuous accumulation model. When the gabor is equiluminant to the background, it has a noisy position signal and the motion dominates the updating of position. From each current location, the next location is determined by extrapolating along the direction of the gabor’s motion. That direction has been shifted by the inappropriate combination of the true external motion and the internal motion so the extrapolated, perceived path drifts farther and farther away orthogonally from the physical path.

Even though this simple model accounts for perceived motion direction, it is not a model of positional mislocalization. In this case, we assume that when position information from the gabor is noisy, as it is on an equiluminant background, position judgments are dominated by predictions based on the gabor’s motion. From each current perceived location, the next location is an extrapolation along the perceived direction (e.g. Nijhawan, 1994; Whitney & Cavanagh, 2000, see Fig. 5). What is unusual here is that the incremental position offsets in the perceived direction continue to accumulate in the direction orthogonal to the physical path, the direction of the internal motion, moving the perceived location further and further from the physical path. The extrapolation in the direction of the gabor’s external motion, simply matches the gabor’s physical displacements in that direction.

One critical factor to generate the double-drift effect is the match between the mean luminance of the gabor and that of the background. With the background maximally different from the gabor mean, as in black background condition of our Experiment 2, the illusion is eliminated. These results suggest that the high degree of positional uncertainty of the gabor on an equiluminant background lets the motion signals contribute to the computation of position. Note that the gabor may have high uncertainty for its position, but the motion of its envelope is registered with sufficient certainty to drive the illusion – with a higher relative weight than the internal motion. With the black background, the positional certainty of the grating patch is sufficient that there is little or no contribution of the internal motion in the computation of location – the value of ***k*** in Equation 2 is 0 when the background is black. Gurnsey and Biard (2012) reported a similar loss of the illusion when they kept the backgound mid-grey but rectified the gabor, replacing the dark bars with the same background grey and leaving only the light bars. Again, with a loss of the match in the mean luminance, the gabor was seen to follow its actual path indicating that its motion was governed only by the external motion and the internal motion was ignored.

Kwon et al. (2015) proposed a tracking model for the perceived deviations of the gabor’s path the integration of the gabor’s motion would contribute to its perceived location when the gabor’s positional uncertainty was low. Their model is similar to our accumulation model above except that they have a saturation of the orthogonal offset that we do not observe in our stimulus conditions. They also proposed that the internal motion that is erroneously attributed to the gabor’s external motion would also be lost from the perceived speed of the internal drift. In other words, if all the internal motion was attributed to the external direction (***k*** = 1), the gabor should appear to have no internal drift. We did not evaluate the apparent velocity of the internal motion but according to their proposal, it should be about 1/5 of the actual speed, since our value of ***k*** was about 0.8, indicating that about 80% of the internal motion was attributed to the external motion of the gabor. Movie 1 can allow an informal judgment of whether this degree of slowing is seen. Its actual internal speed is 5 Hz but Kwon et al.’s model predicts it should appear to be about 1 Hz.

An alternative source for the illusory direction may lie in the paths of the individual features of the gabors (Figure 6). This alternative predicts the same direction deviations and allows some additional inferences about the effects of eccentricity and gabor size. Specifically, the light or dark bars of the gabor themselves travel physically along the direction defined by the two motion vectors. If the gabor were extended to fill the screen as a simple drifting grating, motion selective cells would signal the direction orthogonal to the grating bars – the internal motion direction. However, in the standard gabor, the bars of the grating are short and somewhat blob-like. If the gabor’s bars are short enough, some cells may respond to the oblique displacement of the light or dark feature and this motion response is in the direction that combines the internal and external motions. If this is correct, the illusion should be reduced for gabors with more bars as then there will be competing light and dark segments falling in the same receptive field, cancelling out the motion response. Moreover, the receptive fields of the responding cells need to be large enough to capture the diagonal traversal of a luminance peak or trough predicting that the illusion strength should increase with eccentricity where receptive fields are larger, and decrease for larger gabors. This is indeed what Gurnsey and Biard (2012) report. However, recall that these local motion responses in the illusory direction would still be given by neurons that fall along the actual path; there is no deviation in the position of the stimulus from is actual location. The recovery of the combined vector from the direction of the gabor features cannot alone account for the shift in perceived position that accompanies the shift in direction (Lisi & Cavanagh, 2015).

**Figure 6.**
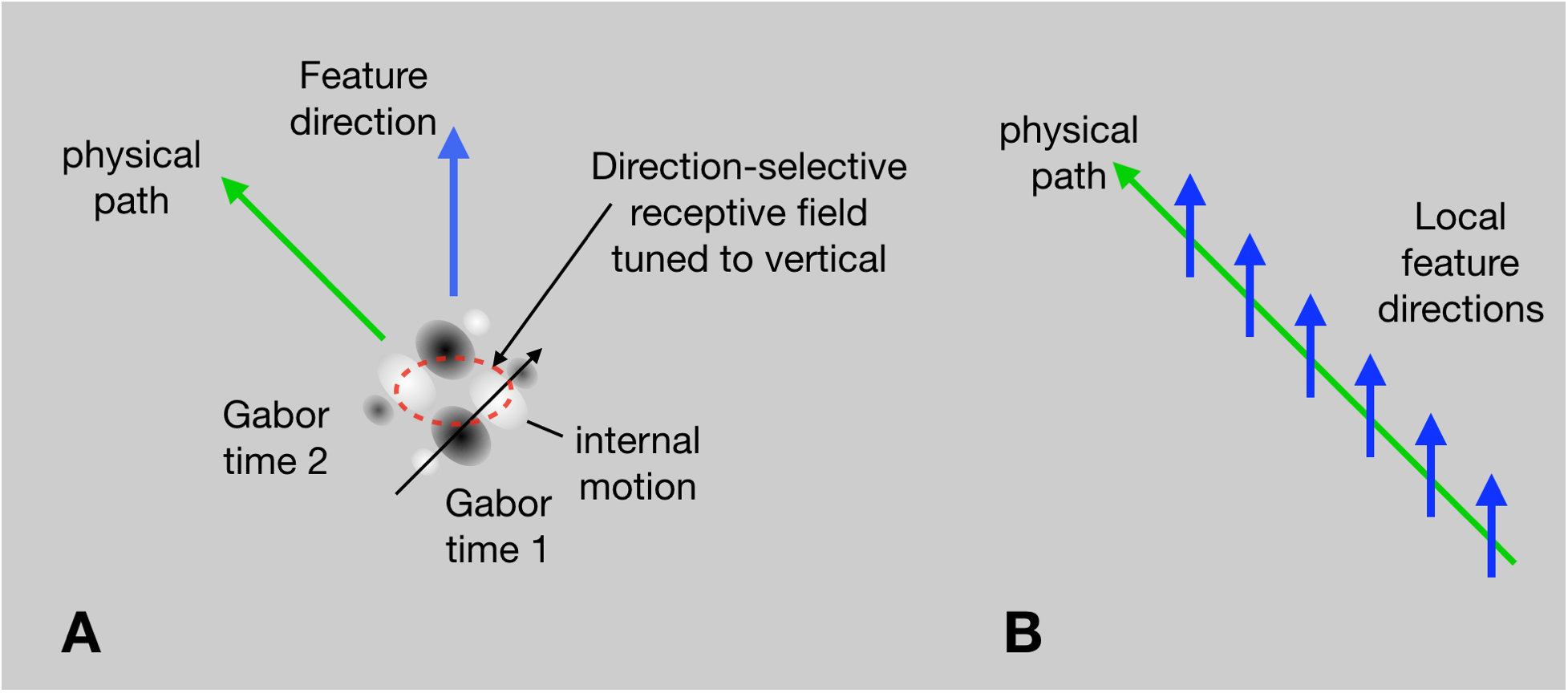
Local feature model for illusory direction. In **A**, the schematized gabor moves up and to the left and its internal motion moves up to the right. The individual light and dark bars within the gabor, however, are drifting straight up (shown as blobs for convenience). The actual path of individual light or dark bars follows the vector combination of the external and internal motion and pass through motion selective receptive fields (red ellipse) that signal that direction. In **B**, the individual features make a series of short upward motions along the physical path as the gabor sweeps up to the left. This is a space-time version of spatial effects like the Frasier spiral where a set of local segments oriented one way are arrayed in a series oriented another way (Morgan, 2015). Note also on the right that the local motions are all oriented upward but there is no position shift. That must arise from accumulating shift that updates position at each moment based on motion direction.

One remarkable property of the double-drift illusion is the duration over which the illusion accumulates, a second or more. This is a striking increase over the 100 ms or so of accumulating position shift seen for a static gabor with moving internal texture (Chung et al, 2007). The direction of individual light or dark features of the static gabor is also consistent with the position shift seen there. Nevertheless, the accumulation of position information from the static gabor envelope appears to quickly override the motion cue and stops the position shift. There are other examples of corrective signals arriving with short time delays. For example, when an oblique grating is moving horizontally within a rectangular aperture, its motion is seen initially perpendicular to the orientation of the bars (Lorenceau, Shiffrar, Wells, & Castet, 1993). However, within 200 ms, the horizontal motion of the terminators propagates inward changing the perceived orientation to horizontal. Similarly, MT neurons initially respond to the direction of motion that is perpendicular to a moving line (Pack & Born, 2001), but over a period of about 60 ms, shift their response properties so that they respond to the true motion of the line independent of its orientation, suggesting that the unambiguously moving endpoints of the line quickly generate a veridical motion solution (see also Pack, Gartland, & Born, 2004). In contrast, any such rapid correction seems to be lost for the double-drift illusion. Possibly, with the moving gabor, the envelope is not stabilized long enough to contradict the position shift suggested by the motion. Our results with pursuit indicate that if the motion of the envelope is the key to avoiding a rapid termination of the accumulating position shift, then that critical motion is in world coordinates, not in retinal coordinates. The gabor is roughly stabilized on the retina during the pursuit condition and yet the position shift continues to accumulate.

## Conclusion

The double-drift illusion offers us a radical mismatch between perceived and retinotopic positions and between perceived and retinotopic motion directions. Areas with cells responding to real-world motion (V3A, V6, MST, VIP) are candidates for the recovery of the combined motion vector that occurs without loss during smooth pursuit. The motion of an object in world coordinates is available here, either computed locally, or biased by top-down feedback from higher level areas. However, our data do not address the question of the anatomical site where the combined motion vector then displaces perceived position continuously accumulating position offsets, but, again, it must happen at or after the locus where the combination with eye movement signals occurs.

## Acknowledgments

The research was support by funding from the Department of Psychological and Brain Science, Dartmouth College (P.C.) and National Science Foundation Grant 1632738 (P.T.). The authors would like to thank Emily Zagans for running the participants.

## References

Brenner, E., & van den Berg, A. V. (1994). Judging object velocity during smooth pursuit eye movements. Experimental Brain Research, 99, 316–324. [PubMed]

Brenner, E., & van den Berg, A. V. (1996). The special rule of distant structures in perceived object velocity. Vision Research, 36, 3805–3814. [PubMed]

Bridgeman, B., Deubel, H., & Haarmeier, T. (1999). Perception and oculomotor behavior in a patient who cannot compensate for eye movements. IOVS, 40, 380.

Cavanagh, P., & Anstis, S. (2013). The flash grab effect. Vision Research, 91, 8–20. https://doi.org/10.1016/j.visres.2013.07.007

Chung, S. T. L., Patel, S. S., Bedell, H. E., & Yilmaz, O. (2007). Spatial and temporal properties of the illusory motion-induced position shift for drifting stimuli. Vision Research, 47, 231–243.

De Valois, R. L. & De Valois, K. K. (1991). Vernier acuity with stationary moving Gabors. Vision Research, 31(9), 1619–1626.

Duhamel, J.R., Bremmer, F., BenHamed, S., Graf, W., 1997. Spatial invariance of visual receptive fields in parietal cortex neurons. Nature 389, 845–848.

Dukelow, S. P., DeSouza, J. F., Culham, J. C., van den Berg, A. V., Menon, R. S., & Vilis, T. (2001). Distinguishing subregions of the human MT+ complex using visual fields and pursuit eye movements. Journal of Neurophysiology, 86(4), 1991–2000.

Eagleman, D. M. & Sejnowski, T. J. (2007). Motion signals bias localization judgments: A unified explanation for the flash-lag, flash-drag, flash-jump, and Frohlich illusions. Journal of Vision, 7:3.

Erickson, R., & Thier, P. (1991). A neuronal correlate of spatial stability during periods of self-induced visual motion. Experimental Brain Research, 86, 608–616.

Fischer, E., Bülthoff, H.H., Logothetis, N.K., & Bartels, A. (2012). Human areas V3A and V6 compensate for self-induced planar visual motion. Neuron 73, 1228–1240.

Freeman, T. C. (2001). Transducer models of head-centred motion perception. Vision Research, 41, 2741–2755. [PubMed]

Freeman, T. C. A., Champion, R. A., Sumnall, J. H., & Snowden, R. J. (2009). Do we have direct access to retinal image motion during smooth pursuit eye movements? Journal of Vision, 9(1):33, 1–11, http://journalofvision.org/9/1/33/, doi:10.1167/9.1.33.

Galletti, C. & Fattori, P. (2003). Neuronal mechanisms for detection of motion in the field of view. Neuropsychologia 41, 1717–1727.

Galletti, C., Squatrito, S., Battaglini, P.P., & Grazia Maioli, M. (1984). ‘Real-motion’ cells in the primary visual cortex of macaque monkeys. Brain Research, 301, 95–110.

Gurnsey, R., & Biard, M. (2012). Eccentricity dependence of the curveball illusion. Canadian Journal of Experimental Psychology, 66(2), 144–152.

Ilg, U. J. & Their, P. (2003). Visual tracking neurons in primate area MST are activated by smooth-pursuit eye movements of an “imaginary” target. Journal of Neurophysiology, 90, 1489–1502.

Ilg, U.J., Schumann, S., & Thier, P. (2004). Posterior parietal cortex neurons encode target motion in world-centered coordinates. Neuron 43, 145–151.

Inaba, N., Shinomoto, S., Yamane, S., Takemura, A., & Kawano, K. (2007). MST neurons code for visual motion in space independent of pursuit eye movements. Journal of Neurophysiology, 97, 3473–3483.

Kwon, O. S., Tadin, D. & Knill, D. C. Unifying account of visual motion and position perception. Proceedings of the National Academy of Sciences of the United States of America 112, 8142–8147, https://doi.org/10.1073/pnas.1500361112 (2015).

Lisi, M., & Cavanagh, P. (2015). Dissociation between the perceptual and saccadic localization of moving objects. Current Biology, 25, 2535–2540.

Lisi, M., & Cavanagh, P. (2017). Different spatial representations guide eye and hand movements. Journal of Vision, 17(2):12. doi: 10.1167/17.2.12

Lorenceau, J., Shiffrar, M., Wells, N., & Castet, E. (1993). Different motion sensitive units are involved in recovering the direction of moving lines. Vision Research, 33, 1207–1217.

Massendari, D., Lisi, M., Collins, T., & Cavanagh, P. (2018). Memory-guided saccades show effect of perceptual illusion whereas visually-guided saccades do not. Journal of Neurophysiology, 119, 62–72. doi: 10.1152/jn.00229.2017

Nau, M., Schindler, A., & Bartels, A. (2018). Real-motion signals in human early visual cortex. Neuroimage, 175, 379–387.

Nijhawan, R. (1994). Motion extrapolation in catching. Nature, 370(6487), 256–257. https://doi.org/10.1038/370256b0

Pack, C. C. & Born, R. T. (2001). Temporal dynamics of a neural solution to the aperture problem in visual area MT of macaque brain. Nature, 22;409(6823):1040–1042.

Pack, C. C., Gartland, A. J., & Born, R. T. (2004). Integration of contour and terminator signals in visual area MT of alert macaque. The Journal of Neuroscience, 24(13), 3268–3280.

Ramachandran, V. S. & Anstis, S. M. (1990). Illusory displacement of equiluminous kinetic edges. Perception, 19(5), 611–616.

Rieger, J. H., & Lawton, D. T. (1985). Processing differential image motion. Journal of the Optical Society of America A, Optics and Image Science, 2, 354–360. [PubMed]

Ross, J., Morrone, M. C., Goldberg, M. E., & Burr, D. C. (2001). Changes in visual perception at the time of saccades. Trends in Neuroscience, 24, 113–121.

Schindler, A. & Bartels, A. (2018). Human V6 integrates visual and extra-retinal cues during head-induced gaze shifts, iScience, 7, 191–197. doi:/10.1016/j.isci.2018.09.004

Shapiro, A., Lu, Z.-L., Huang, C.-B., Knight, E., & Ennis, R. (2010). Transitions between central and peripheral vision create spatial/temporal distortions: A hypothesis concerning the perceived break of the curveball. PLoS One, 5(10), e13296.

Souman, J. L., Hooge, I. T., & Wertheim, A. H. (2005). Perceived motion direction during smooth pursuit eye movements. Experimental Brain Research, 164, 376–386.

Thier, P. & Erickson, R. G. (1992). Responses of Visual-Tracking Neurons from Cortical Area MST-I to Visual, Eye and Head Motion. European Journal of Neuroscience, 4(6), 539–553. PubMed PMID: 12106340.

Tse, P. U. & Hsieh, P. J. J. (2006). The infinite regress illusion reveals faulty integration of local and global motion signals. Vision Research, 46(22), 3881–3885.

Wallach, H. (1959). The perception of motion. Scientific American, 201, 56–60. [PubMed]

Whitney, D., & Cavanagh, P. (2000). Motion distorts visual space: shifting the perceived position of remote stationary objects. Nature Neuroscience, 3(9), 954–959. https://doi.org/10.1038/78878

